# Modulation of ERK1/MAPK3 potentiates ERK nuclear signalling, facilitates neuronal cell survival and improves memory in mouse models of neurodegenerative disorders

**DOI:** 10.1101/496141

**Authors:** Marzia Indrigo, Ilaria Morella, Daniel Orellana, Raffaele d’Isa, Alessandro Papale, Riccardo Parra, Antonia Gurgone, Daniela Lecca, Anna Cavaccini, Cezar M. Tigaret, Gian Michele Ratto, Anna R. Carta, Maurizio Giustetto, Silvia Middei, Raffaella Tonini, Jeremy Hall, Simon Brooks, Kerrie Thomas, Riccardo Brambilla, Stefania Fasano

## Abstract

Cell signalling mechanisms are central to neuronal activity and their dysregulation may lead to neurodegenerative processes and associated cognitive decline. So far, a major effort has been directed toward the dissection of disease specific pathways with the still unmet promise to develop precision medicine strategies. With a different approach, here we show that a selective genetic potentiation of neuronal ERK signalling prevents cell death *in vitro* and *in vivo* in the mouse brain while ERK attenuation does the opposite. This neuroprotective effect can also be induced pharmacologically by a cell permeable peptide mimicking the loss of ERK1 MAP kinase, leading to a selective enhancement of ERK2 mediated nuclear cell signalling. The drug treatment prevents neurodegeneration in mouse models of Huntington’s (HD), Alzheimer’s (AD), and Parkinson’s disease (PD). Importantly, the selective potentiation of ERK2 signalling facilitates both structural and synaptic plasticity, enhances cognition in healthy mice and rescues mild cognitive impairments in both models of AD and HD. Altogether, our observation truly represents a remarkable example of a shared molecular mechanism across multiple neurodegenerative disorders and a potentially valuable therapeutic target for neuro-enhancement.

## Results

The Ras-Raf-MEK-ERK signalling pathway plays a central role in a variety of brain functions. Pharmacological evidence using MEK inhibitors has shown that blockade of both ERK1 and ERK2, the two major isoforms, impairs long-term memory formation and alters other forms of behavioural plasticity ^1–3^. However, previous work indicates that ERK1 and ERK2 play non-redundant roles. Differently from ERK2 KO mice, which are embryonic lethal, ERK1 KO mice showed subtle phenotypes, most notably memory improvements, which are consistent with a repressing role of ERK1 on ERK2 ^4,5^. At the cellular level, the unique N-terminal domain of ERK1 appears to be responsible for the differences in the nuclear localization and the effects on cell proliferation and neuronal cell signaling of the two isoforms ^6,7^.

ERK1 and ERK2 involvement in cell survival remains controversial since both pro-survival and pro-apoptotic mechanisms have been suggested in a number of cell types to explain the effect of MEK inhibition. Moreover, the role of ERK1/2 has not been fully explored *in vivo*, particularly in the context of neurodegenerative disorders. In order to clarify their role in cell survival, we first used selective gene knockdowns of either ERK1 or ERK2 in proliferating mouse embryo fibroblasts (MEF) cells. Lentiviral vectors (LV) either expressing shRNA or shRNAmir for each isoform ^8^ caused opposite effects on cell survival, i.e. reducing ERK1 levels promoted cell survival while ERK2 gene knockdown led to increased apoptosis (Fig. 1a). To confirm these results we predicted that the overexpression of ERK1 would promote cell death by attenuating ERK2 dependent activity. Indeed, we found that the overexpression of ERK1 promoted cell death. Importantly, a similar effect was observed with the overexpression of a chimeric ERK1-like kinase in which the unique N-terminal domain of ERK1 had been fused to ERK2 (hereinafter ERK2>1). In contrast, overexpression of ERK2 or of an ERK2-like kinase (hereinafter ERK1>2), in which the N-terminus of ERK1 had been removed, had little effect on cell survival compared to control cells (Fig. 1b). Consistently, we confirmed these findings in neuronal embryonic cultures where a pro-survival effect induced by ERK1 knockdown and a pro-apoptotic effect induced by ERK2 knockdown was observed (Fig. 1c).

**Figure 1.**
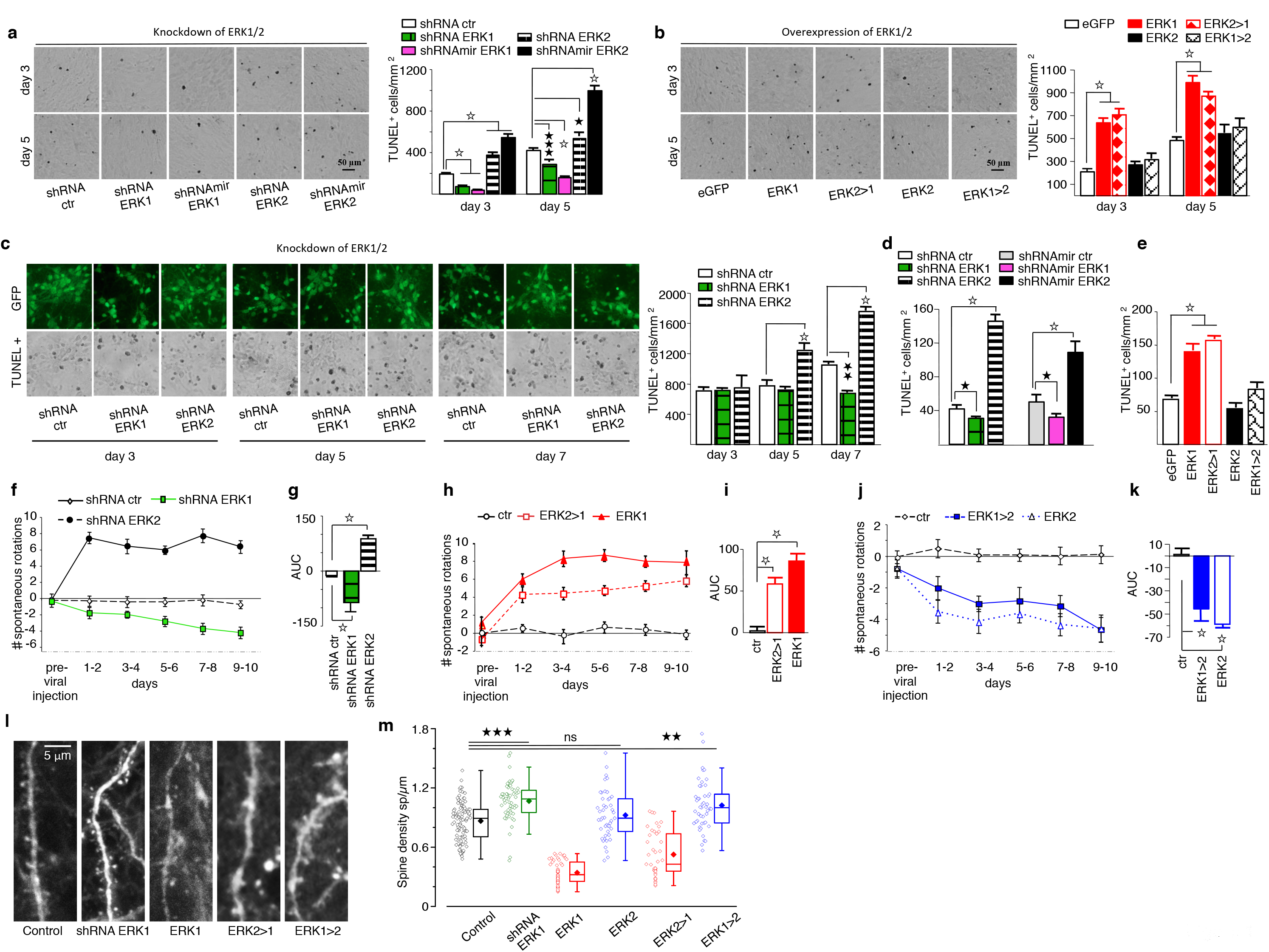
ERK1 MAP Kinase levels negatively regulate ERK2 mediated cell survival. **a** (Left panel) Representative images of TUNEL assay in MEF infected with LV expressing shRNA or shRNAmir for ERK1 or ERK2 at day 3 (top row) and day 5 (bottom row) postinfection. (Right panel) Decreased apoptosis was observed in shRNA ERK1 and shRNAmir-ERK1 infected plates compared to shRNA control plates at day 3 (One-way ANOVA, F_4,80_=178.493 P<0.0001, Bonferroni’s post-hoc, shRNA ctr (n=17) vs shRNA ERK1 (n=18), vs shRNAmir-ERK1 (n=18) P<0.0001) and at day 5 (One-way ANOVA, F_4,107_=281.083 p<0.0001, Bonferroni’s post-hoc, shRNA ctr (n=25) vs shRNA ERK1 (n=27), vs shRNAmir-ERK1 (n=25) p<0.0001). In contrast, a significant increase in apoptotic levels was observed in plates infected with shRNA or shRNAmir for ERK2 at day 3 (Bonferroni’s post-hoc, shRNActr (n=17) vs shRNA ERK2 (n=14), vs shRNAmir-ERK2 (n=18) P<0.0001) and at day 5 (Bonferroni’s post-hoc, shRNA ctr (n=25) vs shRNA ERK2 (n=17) P<0.05, vs shRNAmir-ERK2 (n=18) p<0.0001). **b** (Left panel) Representative images of TUNEL assay in MEF infected with LV overexpressing ERK1, ERK2 and their chimeric constructs. (Right panel) Overexpression of ERK1 or ERK2>1 increased apoptotic levels over eGFP control at day 3 (One-way ANOVA, F_4,45_=80.188 P<0.0001, Bonferroni’s post-hoc, eGFP (n=10) vs ERK1 (n=10), vs ERK2>1 (n=10) P<0.0001) and at day 5 (One-way ANOVA, F_4,50_=53.009 p<0.0001, Bonferroni’s post-hoc, eGFP (n=12) vs ERK1 (n=11), vs ERK2>1 (n=9) P<0.0001). However, overexpression of ERK2 or ERK1>2 did not alter cell survival levels compared to controls. **c** (Left panel) Representative images of apoptosis assay in cortical embryonic cultures expressing shRNA for ERK1 or ERK2 at 3, 5 and 7 days after infection. (Right panel) A significant increase of cell death was observed in shRNA ERK2 group at day 5 (One-way ANOVA, F_2,15_=29.551 P<0.0001, Bonferroni’s post-hoc, shRNA ctr (n=6) vs shRNA ERK2 (n=6) p<0.0001) and at day 7 (One-way ANOVA, F_2,15_=77.076 P<0.0001, Bonferroni’s post-hoc, shRNA ctr (n=6) vs shRNA ERK2 (n=6) P<0.0001). In addition, ShRNA ERK1 diminished apoptotic levels in comparison to control only at day 7 (Bonferroni’s post-hoc, shRNA ctr (n=6) vs shRNA ERK1 (n=6) P<0.01). **d** *In vivo* Knockdown of ERK1 and ERK2. TUNEL assay performed on striatal slices of adult mice injected with either LV expressing shRNA or shRNAmir for ERK1 or ERK2. ERK2 knockdown significantly enhanced cell death while ERK1 knockdown slightly reduced levels of apoptosis in comparison to controls (left panel shRNA, One-way ANOVA, F_2,15_=314.541 P<0.0001, Bonferroni’s post-hoc, shRNA ctr (n=6) vs shRNA ERK2 (n=6) P<0.0001), vs shRNA ERK1 (n=6) P<0.05) (right panel shRNAmir, One-way ANOVA, F_2,15_=57.596 P<0.0001, Bonferroni’s post-hoc, shRNAmir-ctr (n=6) vs shRNAmir-ERK2 (n=6) P<0.0001, vs shRNAmir-ERK1 (n=6) P<0.05). **e** *In vivo* Overexpression of ERK1 and ERK2. TUNEL assay performed on striatal slices of adult mice injected with either LV overexpression of ERK1 or ERK2 and ERK1>2 or ERK2>1. Striatal overexpression of ERK1 or ERK2>1 showed enhanced apoptosis compared to eGFP controls (One-way ANOVA, F_4,15_=28.104 P<0.0001, Bonferroni’s post-hoc, eGFP (n=4) vs ERK1 (n=4) p<0.0001, vs ERK2>1 (n=4) P<0.0001). In contrast, mice with overexpression of ERK2 or ERK1>2 showed similar levels of apoptosis compared to eGFP controls. **f-k** Spontaneous rotational behaviour measured after striatal injection of LV expressing knockdown or overexpression of ERK1 and ERK2 or their chimeric constructs. **f** Control mice (bilateral injection with shRNA ctr) showed a balanced turning behaviour (equal number of 180^0^ rotations either side). Mice with unilateral expression of shRNA ERK1 showed net contralateral rotations (negative values) whereas mice expressing shRNA ERK2 showed increased ipsilateral rotations (positive values) toward the injected side (Repeated Measures Two-way ANOVA, time x virus interaction F_8.497, 276 163_=10.91 1 P<0.0001, ε=.850; viruses effects through time Bonferroni’s post hoc, shRNA ctr (n=26) vs shRNA ERK2 (n=13) at each day P<0.0001; shRNA ctr vs shRNA ERK1 (n=29) day 5-6 P<.05, day 7-8 P<.01, day 9-10 P<0.0001). **g** Analysis of area under curve (AUC) of rotations measured over 10 days showed the cumulative effect caused by ERK1 or ERK2 reduction (Welch’s ANOVA F_2,12_._411_=491.540 P<0.0001, Games-Howell’s post hoc, shRNA ctr vs shRNA ERK1 P<0.0001, shRNA ctr vs shRNA ERK2 P<0.0001). **h** Mice with unilateral overexpression of ERK1 or ERK2>1 showed increased ipsilateral rotations (negative values) toward injection’s side (Repeated Measures Two-way ANOVA, time x virus interaction F_10,190_=8.362 P<0.0001, viruses effects through time Bonferroni’s post hoc, ctr (n=14) vs ERK1 (n=13) at each day P<0.0001, ctr vs ERK2>1 (n=14) day 1-2 P<0.01, from day 3 to day 10 P<0.0001). **i** Analysis of AUC of rotations showed a significant increase of ipsilateral turns compared to control mice (One-way ANOVA, virus effect F_2,24_=312.600 P<0.0001, Bonferroni’s post hoc, ctr vs ERK2>1, vs ERK1 P<0.0001). **j** Mice overexpressing ERK2 or ERK1>2 showed net contralateral rotations (negative values) than control mice (Repeated Measures Two-way ANOVA, time x virus interaction F_10190_=2.832 P<0.01, viruses effects through time Bonferroni’s post hoc, ctr (n=13) vs ERK1>2 (n=14) day 1-2 P<0.05, day 3-4 P<0.01, day 5-6 P<0.05, day 7-8 P<0.01, day 9 10 P<0.0001, ctr vs ERK2(n=14) day 1-2 P<0.01, day 3-4 P<0.0001, day 5-6 P<0.01, from day 7 to day 10 P<0.0001). **k** Analysis of AUC indicated a significant decrease of ipsiateral rotations in comparison with control mice (One-way ANOVA, virus effect, F_2,24_=181.727 P<0.0001 Bonferroni’s post hoc, ctr vs ERK1>2 P<0.0001 and ctr vs ERK2 P<0.0001). **l** Representative images of dendritic spines on striatal neurons after knockdown or overexpression of ERK1 or ERK2. m Mice overexpressing ERK1 or ERK2>1 showed decreased spine density while mice with ERK1 downregulated or overexpressing ERK1>2 had more spines in the striatum (Kolmogorov-Smirnov Test Control vs shRNA ERK1 P<0.001; Control vs ERK1 P<0.001; Control vs ERK2 P=0.4; Control vs ERK1>2 P<0.001; Control vs ERK2>1 P<0.01. Control group (n=9 mice, 105 dendrites) shRNA ERK1 (n=3, 56) ERK1 (n=3, 51) ERK2 (n=3, 46) ERK1>2 (n=3, 47) ERK2>1 (n=3, 40)). Results show mean±s.e.m. White star P<0.0001, ★★★P<0.001 ★★P<0.01, ★ P<0.05.

Next, we tested whether the LV ERK constructs show similar effects also *in vivo*. LV expressing shRNA and shRNAmir targeting ERK1 and ERK2 were injected bilaterally into the dorsal striatum of adult C57Bl/6 mice and two weeks later apoptotic cells expressing GFP fused proteins were counted. ERK2 knockdown promoted cell death while ERK1 ablation slightly reduced basal levels of apoptosis (Fig. 1d). The pro-apoptotic effect of ERK2 ablation *in vivo* was further confirmed by enhanced levels of cleaved caspase-3 (data not shown). Similarly, we observed a significant increase of apoptotic cells in the dorsal striatum of mice bearing the overexpression of ERK1 or the chimeric ERK1-like ERK2>1. On the contrary, overexpression of ERK2 or the chimeric ERK2-like ERK1>2 did not cause significant effects, as similarly observed in neuronal cultures (Fig. 1e).

The striatum is a key structure of the basal ganglia circuits involved in motor control and its dysfunction is associated to a number of neurodegenerative disorders, most notably Parkinson’s and Huntington’s disease. Thus, we explored the possibility that a unilateral alteration of striatal activity mediated by the above ERK1/2 genetic manipulations may impact on spontaneous rotations. Unilateral reduction of striatal ERK1 levels increased contralateral rotations, while reduction of ERK2 induced ipsilateral rotations (Fig. 1f-g). The contralateral sides were injected with control LV to minimise the effect of surgery. Indeed, mice injected with the control LV in both striata showed unbiased turning behaviour. Interestingly, unilateral overexpression of ERK1 and chimeric ERK2>1 increased ipsilateral rotations (Fig. 1h-i), while unilateral overexpression of ERK2 and chimeric ERK1>2 induced contralateral rotations (Fig. 1j-k). These behavioural differences are likely caused by asymmetric alterations of total ERK activity in the striatum. More specifically, the decrease in the ERK2/ERK1 ratio and the concomitant increase in striatal apoptosis in the injected site explain the ipsilateral turning behaviour. To explain the contralateral turning behaviour observed when the ERK2/ERK1 ratio is increased, we speculated that the corresponding change in overall ERK signalling in the injected striatum may enhance structural plasticity and spine formation, two processes highly dependent on ERK activity ^9,10^. Accordingly, analysis of striata injected with the above LVs indicated that altering ERK2/ERK1 ratio resulted in spine density changes on the spiny projection neurons (SPNs). Specifically, while a reduction of ERK1 with shRNA ERK1 increased spine formation, an overexpression of ERK1 (or the chimeric ERK2>1) had the opposite effect. Interestingly, overexpression of ERK2 (or the chimeric ERK1>2) showed a clear trend toward spine density increase (Fig. 1l-m). Together, these results demonstrate that a reduced ERK2/ERK1 ratio is detrimental for cell survival and spine stability raising the hypothesis that a specific potentiation of ERK2-mediated gene regulation could have important therapeutic implications on neuropathological conditions.

The N-terminal sequence of ERK1 appears to be crucial for the observed phenotypes. We previously showed that this sequence is responsible for an altered kinetic of nuclear-cytoplasmic shuttling ^7^. We therefore designed a cell penetrating peptide, hereafter named RB5, corresponding to the 7-38 amino acid sequence of Human MAPK3 (ERK1) aimed to interfere with ERK1 function. In acute striatal slices pre-treated with increasing concentrations of RB5 (0.5-100μM, 1h) we found that RB5 was able to selectively promote ERK activation (Fig. 2a), with an estimated EC50 of 3.4μM. Interestingly, in comparison to a scramble inactive control peptide, RB5 treatment in slices potentiated the phosphorylation of a nuclear target of ERK, histone AcH3 in response to glutamate and blunted the phosphorylation of ribosomal S6 (Thr235/236) protein in the cytoplasm (Fig. 2b-c). In order to examine whether the effect of RB5 may target ERK1 protein to impair its functional contribution to ERK signalling, we repeated the pAcH3 and pS6 assay in ERK1 KO slices. The enhancing effect of RB5 on pAcH3 was indeed mimicked in the absence of ERK1 together with the inhibitory effect on cytoplasmic S6 protein (Fig. 2d-e).

**Figure 2.**
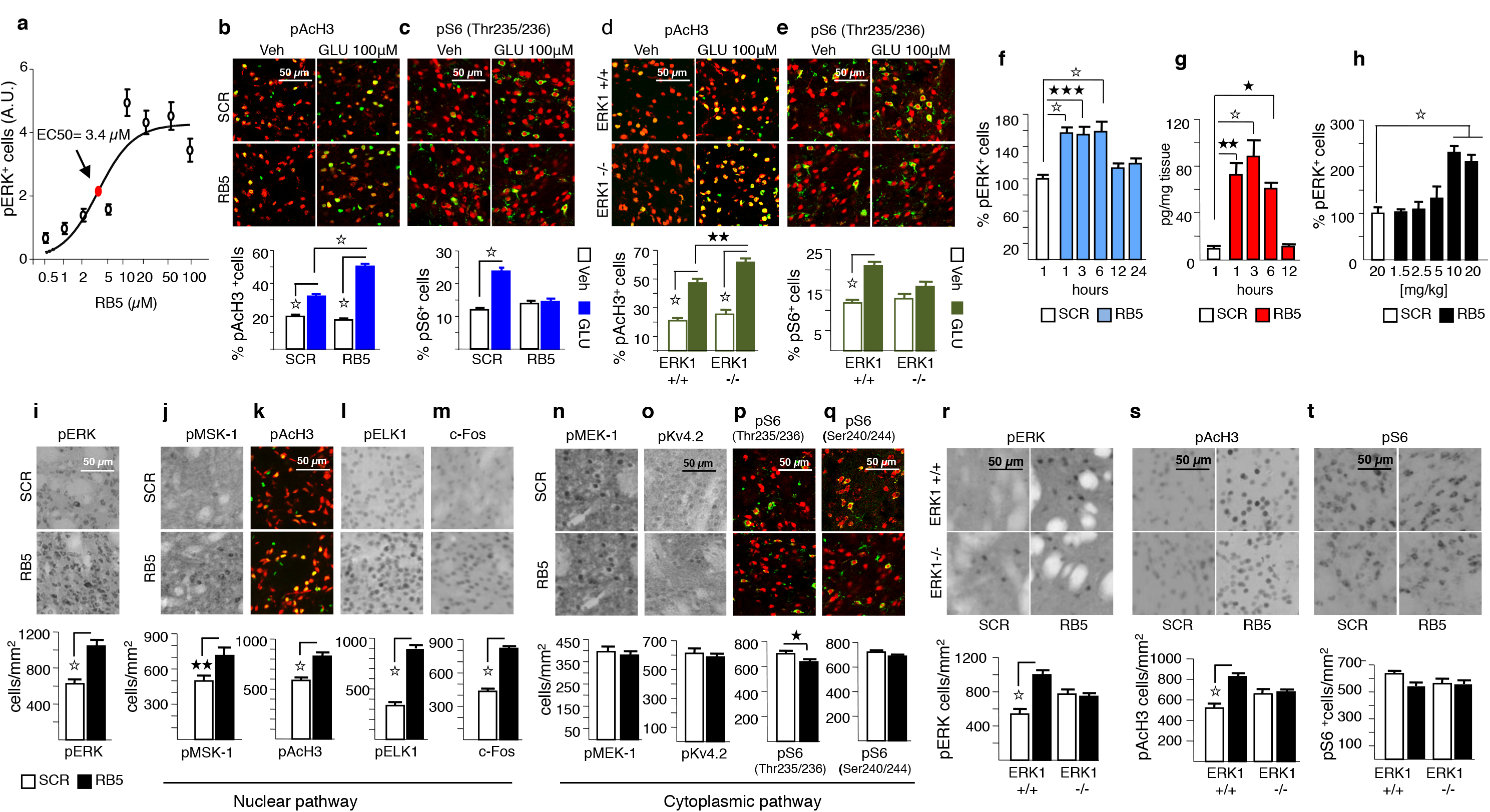
RB5 peptide selectively enhances nuclear ERK signalling. **a** Dose-response curve of RB5 peptide for ERK activation. Different doses of RB5 were used on fresh striatal slices from adult mice. The level of activation is expressed on the Y-axis as arbitrary units (AU). Doses are reported in a logarithmic scale (Log10) on the X-axis. RB5 was effective in enhancing pERK with an EC50 of 3.4 μM. **b-c** (Upper panels) Representative merged images showing histone AcH3 phosphorylation (b) and ribosomal S6 protein phosphorylation (c) on ex vivo acute slices pre-treated for 1h with Scramble (SCR) or RB5 peptide (50μM), and stimulated for 10 min with glutamate (GLU, 100μM) or Vehicle (Veh). (Lower panel b) RB5 enhanced pAcH3 in response to glutamate (Two-way ANOVA, peptide x treatment interaction F_1,59_=73.314 P<0.0001, simple main effect of glutamate F_1,59_=119.285 P<0.0001, Bonferroni’s post-hoc, SCR GLU (n=13) vs RB5 GLU (n=19) P<0.0001). (Lower panel c) RB5 prevented glutamate mediated pS6 increase (Two-way ANOVA, peptide x treatment interaction F_1,45_=40.970 P<0.0001, simple main effect of glutamate F_1,45_=61.369 P<0.0001, Bonferroni’s post-hoc, SCR Veh (n=9) vs SCR GLU (n=12) P<0.0001, RB5 Veh (n=14) vs RB5 GLU (n=14) P=1.000). **d-e** (Upper panels) Representative merged images showing pAcH3 (d), and pS6 (e) on slices from ERK1-/-mice pre-treated for 1h with SCR or RB5 peptide (50μM) and stimulated with glutamate or vehicle for 10 min. (Lower panel d) ERK1-/- slices showed a greater enhancement of pAcH3 in response to glutamate (Two-way ANOVA, genotype x treatment interaction F_1,41_=5.448 P<0.05, simple main effect of glutamate F_1,41_=9.617 P<0.01, Bonferroni’s post-hoc, ERK1-/- GLU (n=11) vs ERK1+/+ GLU (n=11) P<0.01). (Lower panel e) Absence of ERK1 prevented glutamate mediated pS6 enhancement (Two-way ANOVA, genotype x treatment interaction F_1,43_=9.900 P<0.01, simple main effect of glutamate F_1,43_=11.315 P<0.01, Bonferroni’s post-hoc, ERK1+/+ GLU (n=15) vs ERK1-/- GLU (n=8) P<0.01, ERK1-/- Veh (n=9) vs ERK1-/- GLU P=0.701). **f** PERK positive cells in dorsal striatum induced by a single dose of RB5 (20 mg/kg, i.p.) at different time-points (n=5 mice/group). Scramble-injected mice (20 mg/kg, i.p.) were perfused after 1h. RB5-mediated enhancement of ERK phosphorylation lasted up to 6h post-injection (One-way ANOVA, F5,2_4_=10.942 p<0.0001, Bonferroni’s post-hoc, SCR vs RB5 1h and 6h P<0.0001, SCR vs RB5 3h P<0.001). **g** RB5 brain levels measured with mass spectrometry at different hours after a single administration (20 mg/kg i.p.). High levels of RB5 were detected up to 6h (Oneway ANOVA, F_4,18_=17.469 P<0.0001, Bonferroni’s post-hoc, SCR vs RB5 1h P<0.01, SCR vs RB5 3h P<0.0001, SCR vs RB5 6h P<0.05). **h** PERK positive cells in dorsal striatum induced by increasing doses of RB5 (1.5, 2.5, 5, 10, and 20 mg/kg, i.p.) at 1h after administration (n=5 mice/group). SCR peptide was injected at 20 mg/kg, i.p. All mice were perfused after 1h. RB5 caused a significant increase of ERK phosphorylation at both 10 and 20 mg (One-way ANOVA, F_5,24_=14.292 P<0.0001, Bonferroni’s post-hoc, SCR vs RB5 10, and vs RB5 20 P<0.0001). **i-t** (Upper panels) Representative images showing p-ERK, nuclear ERK-dependent markers and cytoplasmic markers in striata of mice injected either with RB5 or SCR peptide (20 mg/kg, i.p.) and perfused 1h later. (Lower panels) RB5 increased ERK phosphorylation (**i**, Independent sample t-test t_28_= −5.340 P<0.0001; SCR (n=15) RB5 (n=15)) and promoted a selective activation of nuclear ERK-dependent markers as pMSK-1 (**j**, Independent sample t-test t_29_= −2.808 P<0.01; SCR (n=16) RB5 (n=15)), pAcH3 (**k**, Independent sample t-test t_28_= −5.404 P<0.0001; SCR (n=16) RB5 (n=14)), pELK (**l**, Independent sample t-test t_29_= −12.106 P<0.0001; SCR (n=16) RB5 (n=15)) and c-Fos (**m**, Independent sample t-test t_29_= −12.743 P<0.0001; SCR (n=16) RB5 (n=15)). Differently, in the cytoplasm RB5 did not affect the phosphorylation of pMEK-1 (**n**, Independent sample t-test t_28_= 0.609 P=0.547; SCR (n=15) RB5 (n=15)) or the phosphorylation of Kv4.2 (**o**, Independent sample t-test t_28_= 0.618 P=0.541; SCR (n=16) RB5 (n=14)). In addition, RB5 slightly reduced ERK dependent phosphorylation of S6 (Thr235/236) (**p**, Independent sample t-test t_29_= 2.191 p<0.05; SCR (n=16) RB5 (n=15)) while had no effect on mTOR/TORC1 dependent phosphorylation of S6 (Thr240/244) (**q**, Independent sample t-test t_28_= 1.878 P=0.071; SCR (n=16) RB5 (n=14)). **r-t** (Upper panels) Representative images of phosphorylation of ERK, AcH3 and S6 in striata of ERK1-/- mice after a single RB5 injection. **r** (Lower panel) RB5 increased ERK phosphorylation in WT but not in KO mice (Two-way ANOVA, genotype x peptide interaction F_1,36_=23.301 P<0.0001, simple main effect of peptide F_1,36_=37.707 p<0.0001, Bonferroni’s post-hoc, ERK1+/+ SCR (n=10) vs ERK1+/+ RB5 (n=8) P<0.0001; ERK1 -/- SCR (n=12) ERK1 -/- RB5 (n=10)). **s** (Lower panel) RB5 increased pAcH3 in WT mice (Two-way ANOVA, genotype x peptide interaction F_1,30_=17.494 P<0.0001, simple main effect of peptide F_1,30_=37.505 P<0.0001, Bonferroni’s post-hoc, ERK1+/+ SCR (n=8) vs ERK1+/+ RB5 (n=8) P<0.0001; ERK1 -/- SCR (n=9) ERK1 -/- RB5 (n=9)). t (Lower panel) RB5 treatment did not change the phosphorylation of S6 (Thr235/236) in both genotypes (Two-way ANOVA, genotype x peptide interaction F_1,31_=2.601 P=0.117, main effect of peptide F_1,31_=3.266 P=0.080, main effect of genotype F_1,31_=1.185 P=0.285; ERK1+/+ SCR (n=9) ERK1+/+ RB5 (n=10) ERK1-/- SCR (n=8) ERK1-/- RB5 (n=8)). Results show mean±s.e.m. White star P<0.0001, ★★★P<0.001, ★★ P<0.01, ★ P<0.05.

In order to use RB5 *in vivo*, we subsequently perfomed a preliminary pharmacokinetic and pharmacodynamic characterisation of the peptide. Upon a single administration of the peptide *in vivo*, ERK phosphorylation remained sustained up to 6h post injection, which suggests a half-life of between 6 and 12h (Fig. 2f). The brain bioavailability of RB5 was confirmed by mass spectrometric analysis which indicated detectable levels of RB5 in most brain areas up to 6h post injection (Fig. 2g). Moreover, in a dose response study, both 10 and 20 mg/kg i.p. resulted in significant ERK activation *in vivo* (Fig. 2h). We acutely administered RB5 or its scamble control (20 mg/kg, i.p.) to mice and 1h later we collected their brains for further analyses with IHC and IF. RB5 greatly enhanced striatal ERK phosphorylation in comparison to the scramble peptide (Fig. 2i) and promoted a selective activation of nuclear ERK-dependent signalling and gene transcription (pMSK, pAcH3, pELK, c-Fos)(Fig. 2j-m). In contrast, RB5 did not influence the phosphorylation of MEK-1 (Fig. 2n) or the phosphorylation of voltage-gated K^+^ (Kv4.2) channels in the cytoplasm, two established targets of ERK (Fig. 2o). Moreover, ERK dependent phosphorylation of S6 (Thr235/236) was also slightly reduced similar to *in vitro* experiments (Fig. 2p). As a confirmation of the selectivity of RB5 action on ERK signalling we confirmed that no effect was observed on TORC1 dependent phosphorylation of S6 (Thr240/244) (Fig. 2q).

Besides the preferential activation of the nuclear pathway, RB5 action seems to resemble what has been previously observed in ERK1-deficient cells in which the stimulus-dependent activation of ERK2 was enhanced ^4^ (see also Fig. 2d-e). To confirm this observation, a single administration of RB5 *in vivo* induced a significant activation of pERK and pAcH3 in WT mice but this effect was lacking in ERK1 KO mice, suggesting that RB5 behaves as an ERK1 inhibitor (Fig. 2r-s). As an additional control we showed that phosphorylation of S6 (Thr235/236) was comparable in both genotypes, confirming that RB5 preferentially activates nuclear signalling (Fig. 2t). Overall, these data demonstrate that RB5, by mimicking the increase of ERK2/ERK1 ratio and by selectively stimulating ERK-mediated gene expression and epigenetic remodelling in the brain, can be considered as a pharmacological model of ERK1 KO/knockdown.

We then examined whether an increase in ERK signalling in the brain could prevent neurodegeneration in models of three pathological conditions: Huntington’s Disease (HD), Parkinson’s Disease (PD) and Alzheimer’s Disease (AD). To address the possibility that ablation of ERK1 and overexpression of ERK2 might play a protective role against neurodegeneration, we first used the neurotoxin 3-nitropropionic acid (3-NP) model of HD ^11,12^. Mice underwent bilateral striatal injection of LV expressing shRNA ERK1 (Fig. 3a), ERK2 or the chimeric ERK1>2 (Fig. 3b) and two weeks later they were administered with 3-NP (50 mg/kg, i.p.) for 21 days. 3-NP-induced striatal degeneration as measured with TUNEL staining was significantly reduced in all groups and comparable with saline treated animals. A parallel experiment in which ERK1 KO mice were chronically administered with 3-NP revealed mostly similar effects of reduced cell death in the striatum (Fig. 3c). Importantly, WT animals co-treated with RB5 (20 mg/kg, i.p. twice a day, 7 days) and 3-NP (50 mg/kg, i.p.) revealed a significant reduction of neuronal death (Fig. 3d-e). In Hdh^Q111/+^ knock-in mice, a late onset genetic model of HD ^13,14^, RB5 (20 mg/kg, i.p. twice a day, 7 days) was able to attenuate striatal neuronal degeneration (Fig. 3f) and restore the ERK enhancing effect in comparison to scramble peptide (Fig. 3g).

**Figure 3.**
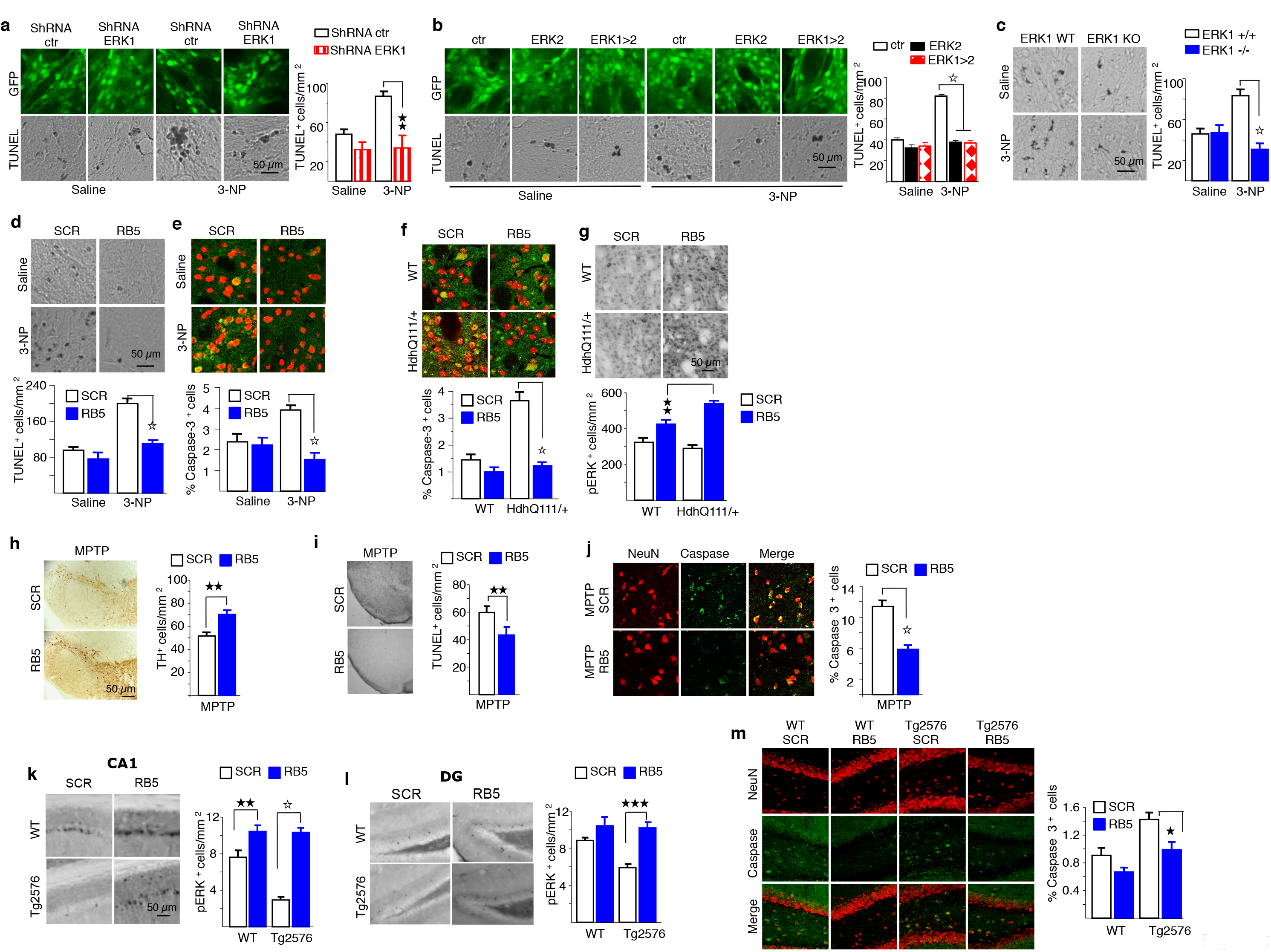
RB5 shows neuroprotective effects in HD, PD and AD mouse models. **a-b** (Left panels) Representative images of TUNEL assay in mice with *in vivo* striatal knockdown of ERK1 (a), overexpression of ERK2 and ERK1>2 (b). (Right panels) TUNEL assay performed after a subchronic treatment of 21 days with either 3-NP (50 mg/kg, i.p. once a day) or saline. ShRNA ERK1 protected striatal neurons from 3-NP induced apoptosis (**a** right panel, Two-way ANOVA, virus x treatment interaction F_1,15_=4.953 P<0.05, simple main effect of virus F_1,15_=21.222 P<.0001, Bonferroni’s post-hoc, shRNA ctr 3-NP (n=5) vs shRNA ERK1 3-NP (n=5) P<0.0001; shRNA ctr Saline (n=4), ERK1 Saline (n=5)) as well as ERK2 and ERK1>2 (**b** right panel, Two-way ANOVA, virus x treatment interaction F_2,33_=66.32O p<0.0001, simple main effect of the viruses F_2,33_=194.003 P<0.0001, Bonferroni’s post-hoc, ctr 3-NP (n=10) vs ERK2 3-NP (n=6), vs ERK1>2 3-NP (n=5) p<0.0001; ctr Saline (n=8), ERK2 Saline (n=5), ERK1>2 Saline (n=5)). **c** (Left panel) Representative images of TUNEL assay in the striatum of ERK1-/- mice injected either with saline or 3-NP (50 mg/kg, i.p. once a day) during 21 days. (Right panel) ERK1-/- mice were protected against the 3-NP induced neurotoxicity in the striatum (Two-way ANOVA, genotype x treatment effect F_1,41_=2O.629 P<0.0001, simple main effect of genotype F_1,41_=44.647 p<0.0001, Bonferroni’s post-hoc, ERK1+/+ 3-NP (n=13) vs ERK1-/- 3-NP (n=12) P<0.0001; ERK1 +/+ saline (n=11), ERK1 -/- Saline (n=9)). **d-e** (Upper panels) Representative images of TUNEL assay and caspase-3 immunofluorescence (NeuN+ in red, caspase 3+ in green) in the striatum of mice injected either with SCR or RB5 peptide (20 mg/kg, i.p., twice a day, 12h interval) and with saline or 3-NP (50mg/kg, i.p., once a day) for 7 consecutive days. (Lower panels) Mice co-treated with RB5 and 3-NP showed reduced apoptotic levels (**d**, Two-way ANOVA, peptide x toxin interaction F_1,33_=13.232 P<0.001, simple main effect of peptide F_1,33_=44.250 p<0.0001, Bonferroni’s post-hoc, SCR 3-NP (n=9) vs RB5 3-NP (n=10) P<0.0001; SCR saline (n=9), RB5 saline (n=9)) and diminished caspase-3 levels (e, Two-way ANOVA, peptide x toxin interaction F_1,26_=12.700 P<0.001, simple main effect of peptide F_1,26_=30.556 p<0.001, Bonferroni’s post-hoc, SCR 3-NP (n=9) vs RB5 3-NP (n=7) P<0.0001; SCR saline (n=7) vs RB5 saline (n=7)). **f-g** (Upper panels) Representative images of Hdh^Q111/+^ caspase-3 and pERK respectively, of Hdh^Q111/+^ mice injected either with RB5 or SCR (20 Q111/+ mg/kg i.p.) for 8 days. (Lower panels) Hdh^Q111/+^ mice treated with RB5 showed diminished caspase-3 levels (**f**, Two-way ANOVA, peptide x genotype interaction F_1,16_=7.125 P<0.05, Q111/+ main effect of peptide F_1,16_=31.354 P<0.0001, Bonferroni’s post-hoc, Hdh^Q111/+^ SCR (n=5) vs Hdh^Q111/+^ RB5 (n=5) P<0.0001; WT SCR (n=5), WT RB5 (n=5)) and a greater enhancement of ERK phosphorylation compared to WT mice (**g**, Two-way ANOVA, peptide x genotype interaction F_1,29_=15.851 P<0.0001, main effect of genotype F_1,29_=18.605 P<0.0001, Bonferroni’s post-hoc, Hdh^Q111/+^ RB5 (n=9) vs Hdh^Q111/+^ SCR (n=9) P<0.0001, WT RB5 (n=8) vs Hdh^Q111/+^ vs Hdh RB5 P<0.01, WT SCR (n=7) vs WT RB5 P<0.01). **h-j** Representative images showing TH, TUNEL and Caspase-3 respectively, after co-treatment with MPTP (20 mg/kg i. p.) and RB5 or SCR (20 mg/kg i.p.) for 4 days. RB5 showed a protective effect on DA cells of the SN from MPTP toxicity (**h**, right panel, Independent sample t-test t_10_= −3.884 p<0.01, MPTP SCR (n=5) MPTP RB5 (n=7)) and a concomitant decrease of apoptotic cells (i, right panel, Independent sample t-test t_9_=-3.551 P<0.01, MPTP SCR (n=5) MPTP RB5 (n=6)) and caspase-3 (**j**, right panel, Independent sample t-test t_9_=6.433 P<0.0001, MPTP SCR (n=5) MPTP RB5 (n=6)). **k-m** Representative images showing pERK in CA1 and DG and caspase-3 in DG of Tg2576 mice treated either with RB5 or SCR (20 mg/kg i.p.) for 7 days. RB5 significantly enhanced ERK phosphorylation in both CA1 (**k**, Two-way ANOVA, peptide x genotype interaction F_1,23_=18.245 P<.0001, simple main effect of peptide F_1,23_=90.240 P<0.0001, Bonferroni’s post-hoc, Tg2576 RB5 (n=8) vs Tg2576 SCR (n=6) P<0.0001, WT RB5 (n=7) vs WT SCR (n=6) P<0.01) and DG (l, Two-way ANOVA, peptide x genotype interaction F_1,23_=12.099 P<.05, simple main effect of peptide F_1,23_=28.431 P<0.0001, Bonferroni’s post-hoc, Tg2576 RB5 (n=7) vs Tg2576 SCR (n=6) P<0.0001; WT SCR (n=7) WT RB5 (n=7)). In addition, Tg2576 mice treated with RB5 showed lower levels of caspase-3 compared to untreated mice (**m**, Two-way ANOVA, simple main effect of peptide F_1,26_=14.088, p<0.001, simple main effect of genotype F_1,26_=21.857, P<0.0001, Bonferroni’s post-hoc, Tg2576 SCR (n=7) vs Tg2576 RB5 (n=7) P<0.05; WT SCR (n=7) vs Tg2576 SCR P<0.01; WT RB5 (n=9)). Results show mean±s.e.m. White star P<0.0001, ★★★P<0.001, ★ ★ P<0.01, ★ P<0.05.

Next, we used the 1-methyl-4-phenyl-1,2,3,6-tetrahydropyridine (MPTP) mouse model of PD to investigate the therapeutic potential of RB5 ^15-17^. Adult mice were pre-treated with RB5 (20 mg/kg i.p.) or scramble peptide before MPTP (20 mg/kg i.p.) once a day for 4 consecutive days. Mice were then killed on day 5 and brain slices were prepared for immunohistochemical analysis. RB5 prevents tyrosine hydroxylase (TH) loss in the SNc (Fig. 3h) and counteracts apoptotic cell death induced by sub-acute MPTP (Fig. 3i-j).

Finally, in Tg2576 transgenic mice, a genetic model of AD, a 7 day RB5 treatment (20 mg/kg, i.p.) was able to restore ERK phosphorylation levels in both the CA1 and the dentate gyrus hippocampal regions (Fig. 3k-l) and to significantly reduce the pre-apoptotic phenotype as measured by caspase-3 (Fig. 3m).

The observed effects described so far indicate that a positive modulation of ERK activity in the brain results in a neuroprotective action across different models of neurodegeneration. However, ERK-dependent signalling also plays a major role in cognition and its selective nuclear potentiation could lead to a cognitive enhancement. Significantly, early work using ERK1 KO mice clearly indicated that ERK2 potentiation leads to enhanced synaptic plasticity and memory ^4,5^ but the evidence has been so far limited to a single mouse model in which ERK1 has been ablated from embryonic development, with the associated confounding developmental effects. Therefore, we tested the hypothesis that a pharmacological potentiation of nuclear ERK signalling in the adult may lead to significant synaptic and behavioral changes, both in WT mice and in models of disease.

To investigate the effect of RB5 on synaptic transmission in the hippocampus we performed whole-cell patch clamp recordings of excitatory synaptic responses in acute slices, at Schaffer collateral (SC)-CA1 synapses, in which ERK signalling plays a prominent role ^18^. Intracellular dialysis of the RB5 peptide (50 μM) induced a gradual but sustained increase in the amplitude of excitatory postsynaptic currents (EPSCs) evoked in CA1 pyramidal neurons by stimulation in *stratum radiatum*, whereas the infusion of the scramble control peptide (50 μM) had no effect (Fig. 4a). This result indicates that RB5 is able to positively modulate synaptic efficacy in a tetanus-independent manner in the hippocampus.

**Figure 4.**
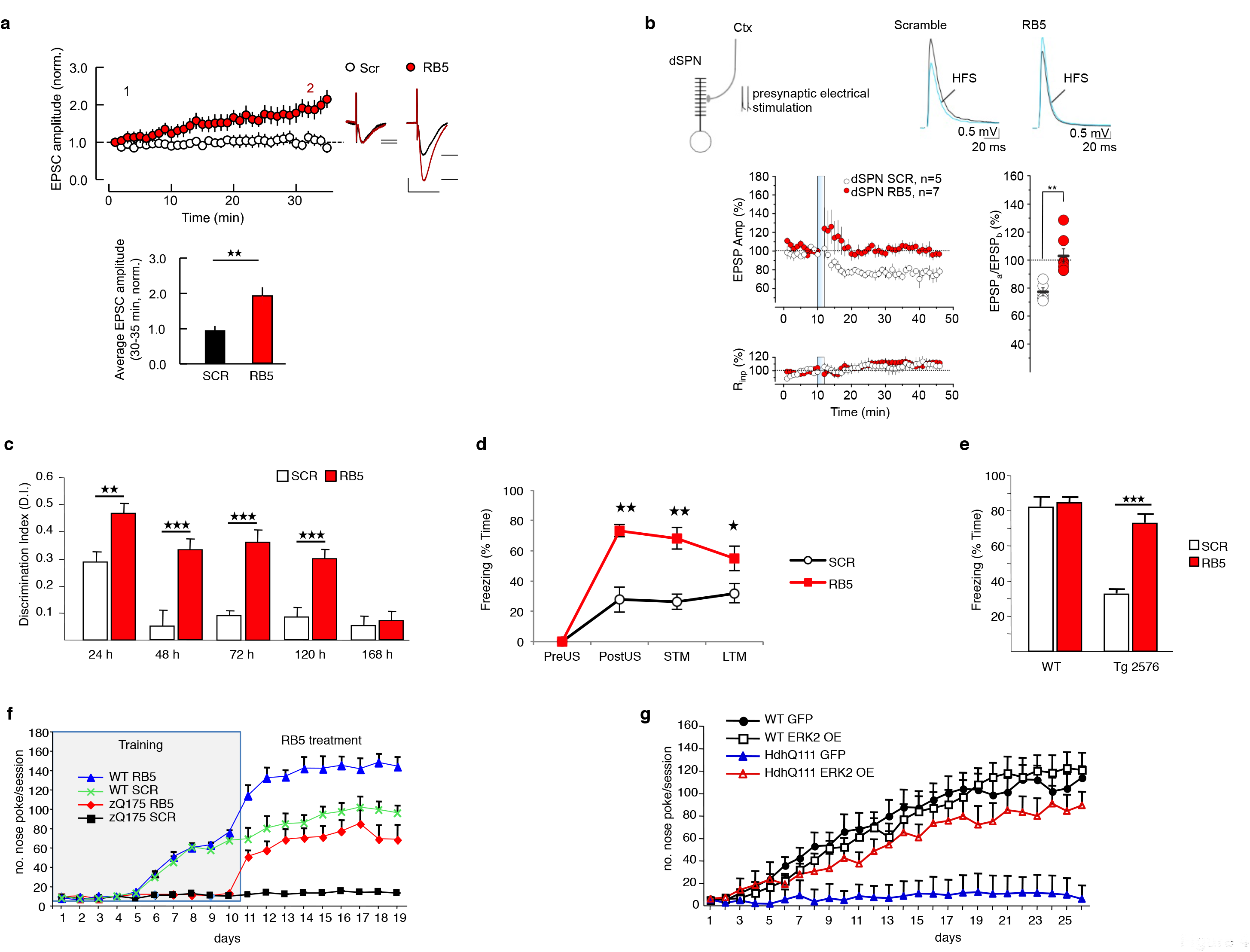
ERK enhancement facilitates both synaptic plasticity and memory and contrasts cognitive deterioration. **a** Intracellular dialysis of RB5 peptide increased evoked synaptic currents at SC-CA1 synapses, whereas the control peptide (SCR) had no effect. (Upper panel) Time course of minute-average EPSC amplitude at SC-CA1 synapses in whole-cell patch clamp recordings with RB5 (50μM) or SCR peptide (50μM) in the pipette solution (values normalized to the mean EPSC amplitude during the first minute of recording). (Insets) Representative current traces from 3 min (black, 1) and 33 min time points (red, 2). (Lower panel) Average EPSC amplitude at 30-35 min, normalized to first minute of recording. Data shown as mean±s.e.m. RB5 (1.928±0.246) (n=12 cells, 4 animals) and SCR (0.938±0.141) (n=8 cells, 4 animals). Independent sample t-test assuming unequal variances t_16,46_= 3.65, P<0.01 confidence interval: 95%. Scale bars: 40pA and 50ms. **b** (Top Left) Schematic representation of electrically stimulated glutamatergic cortical inputs to dSPNs. (Top Right) Superimposed averaged recordings (ten traces) before and after the delivery of the HFS protocol. (Bottom left) The HFS protocol (vertical bar) failed to induce plasticity in dSPNs when the RB5 peptide (50μM) was intracellularly applied (RB5, 103±5 % of baseline, n=7, Repeated Measures Oneway ANOVA F_6,44_= 0.8, P=0.45). The specificity of action of RB5 was confirmed on its control peptide (50μM), which had no effect on plasticity induction (SCR, 77±3 % of baseline, n=5; P<0.05, Repeated Measures One-way ANOVA F_4,44_=4.2 P<0.05, Tukey’s post hoc P<0.05; dSPN_RB5 vs dSPN_SCR, Mann Whitney test P<0.01). Data are shown as a time-course (mean±s.e.m.) of normalized EPSP amplitudes and normalized Rinp. (Bottom right) Scatterplot summarizes the ratios of synaptic responses after (a; 20-30 min) and before (b; 5 min) the HFS. c RB5 significantly enhanced memory formation and consolidation in Novel Object Recognition Test (NOR) up to 5 days. Discrimination Index (D.I.) was tested in independent groups at different time points 24, 48, 72, 120 and 168h. At 24h, D.I. of RB5 group was significantly higher than the one of the SCR group (Independent sample t-test t_26_= 4.428 P<0.01, SCR (n=14) RB5 (=14)) as it was at 48h (Independent sample t-test t38= 4.210 P<0.0001, SCR (n=18) RB5 (=22)). At 48h D.I. of Scramble group dropped below chance level (One sample t-test SCR t_17_= 0.566 P=0.579) while D.I. of RB5 group was greatly above it (One sample t-test RB5 t_21_= 8.077 P<0.0001). At 72h, D.I. of RB5 group remained significantly higher than the one of SCR (Independent sample t-test t_14_= 7.899 P<0.0001, SCR (n=7) RB5 (=9)) and above chance level (One sample t-test RB5 t_6_= 12.341 P<0.0001; SCR t_6_= 9.073 P<0.0001). At 120h, D.I. of RB5 group was still significantly higher than SCR (Independent sample t-test t_23_= 4.290 P<0.0001, SCR (n=10) RB5 (=15)) and above chance level (One sample t-test RB5 t_14_= 9.143 P<0.0001; SCR t_9_= 2.230 P=0.053). At 168h, D.I. of RB5 group finally went down to chance level, indistinguishable from SCR group (Independent sample t-test t_18_= 0.777 P=0.447, SCR (n=10) RB5 (=10)). Data are expressed as mean±s.e.m. **d** RB5 enhanced acquisition of contextual fear memory in rats. Awake rats were infused bilaterally via indwelling steel cannula aimed at the dorsal CA1 with either 2mg/ml SCR peptide (n=6) or RB5 (n=5) 2Omin prior to conditioning. STM and LTM were assessed by measuring the conditioned fear behaviour during a 2min recall test 3h and 2 days after training, respectively. RB5 administration had a profound effect on the freezing behaviour of the rats (Repeated Measures Two-way ANOVA, test x group interaction, F_3,27_=9.439 p<0.001; group effect, F_1,9_=23.171 P<0.001). Freezing behaviour of RB5 rats was significantly increased, compared with the one of SCR rats, in the PostUS, STM and LTM tests (Post hoc t-tests, PostUS: P<0.01, STM: P<0.01, LTM: P<0.05). **e** RB5 enhanced acquisition of Contextual Fear memory in the Tg2576 mouse model of Alzheimer’s Disease. 7 months old Tg2576 mice and WT mice were treated for 1h with either SCR or RB5 (20 mg/kg, i. p.) before Contextual Fear Conditioning training and tested for memory retention 24h later. Contextual Fear memory impairment in Tg2576 mice was fully rescued by RB5 treatment. Values are reported as % of time spent freezing (Two-way ANOVA genotype x treatment interaction F_1,26_= 19.583 P<0.0001. Bonferroni post-hoc Tg2576 SCR (=7) vs Tg2576 RB5 (n=8) p<0.0001). Data are expressed as mean±s.e.m. **f** RB5 treatment improved performances of both WT and zQ175 mice in the 9-hole operant conditioning task. WT (n=31) and zQ175 (n=34) mice underwent daily 20 minutes nose poke training for 10 consecutive days (days 1-10) on a simple fixed ratio (FR1) schedule of reinforcement. Starting from day 11 mice received one administration of RB5 (10 mg/kg. i.p.) 1h before the nose poke training. RB5 treatment improved performances of both WT and zQ175 female mice (Repeated Measures Three-way ANOVA, time effect F_2.367,61.550_= 8.0 54 P<0.0001, genotype effect F_1,26_=90.156 p<0.0001, treatment effect F_1,26_=45.390 P<0.0001, Bonferroni’s post hoc, zQ175 RB5 vs zQ175 SCR day11-12 P<0.05, from day 13 to day 17 P<0.0001, day 18-19 P<0.01; zQ175 SCR vs WT SCR from day 11 to day 19 P<0.0001; WT SCR vs WT RB5 from day 11 to day 16 P<0.01, day 17 P=0.077, day 18-19 P<0.01). zQ175 RB5 (n=7) zQ175 SCR (n=8) WT RB5 (n=7) WT SCR (n=8). **g** WT (n=19) and Hdh^Q111/+^ mice (n=17) injected at 2 months of age with either LV-GFP or LV-overexpressing ERK2 (ERK2 OE) underwent nose poke training at 18 months of age. Untreated Hdh^Q111/+^ mice significantly differed from the other three groups and poorly performed in this task while Hdh^Q111/+^ treated mice were indistinguishable from the WT controls (Repeated Measures Three-way ANOVA, time effect F_3.309, 105.898_=71.004 p<0.0001; genotype effect F_1,32_=22.684 P<0.0001; treatment effect F_1,32_=6.658 p<0.05, Bonferroni’s post hoc, Hdh^Q111/+^ GFP (n=9) vs HdhQ111 ERK2 OE (n=8) from day 12 to day 16 P<0.05, day 17-18 P<0.01, day 19 P0.05, from day 20 to day 25 p<0.01, day 26 P<0.0001; Hdh^Q111/+^ GFP vs WT GFP (n=10) from day 8 to day 11 P<0.01, from day 12 to day 26 P<0.0001). ★★★P<0.001, ★★ P<0.01, ★ P<0.05.

We next asked whether RB5 could modulate synaptic activity also in the striatum, a brain region in which ERK signalling has been highly implicated in the control of certain forms of long term potentiation (LTP) and long term depression (LTD) ^19,20^. We previously showed that at the corticostriatal synapses on direct spiny projection neurons (dSPNs), high-frequency stimulation (HFS) of the deep cortical layer V of the somatosensory cortex results in a form of adenosine A1 receptor (A1R)-mediated LTD; this LTD is suppressed upon the concurrent activation of postsynaptic dopamine D1R, or downstream signalling cascades, which include the ERK pathway ^20,21^. Thus, we determined whether the direct activation of ERK signalling cascade using RB5 could affect this form of LTD. We found that the stimulation protocol failed to induce LTD when RB5 peptide was administered via a patch pipette to the dSPN, while the scramble peptide did not alter this form of synaptic plasticity (Fig. 4b). Altogether these data support the notion that a selective potentiation of nuclear ERK signalling is sufficient to modify synaptic efficacy in two distinct circuitries of the brain.

The clear effect on synaptic plasticity mediated by RB5 in brain slices prompted us to investigate whether a single administration of this ERK enhancing peptide could improve cognition in normal mice. In the widely used novel object recognition (NOR) test ^22,23^, a single systemic RB5 administration resulted in a sustained memory consolidation (Fig. 4c). Remarkably, the NOR memory remained highly significant up to 120h post training while for the scramble peptide-treated animals the discrimination index returned to chance levels within 48h. NOR learning is largely dependent on the perirhinal cortex whilst declarative like memory is dependent on the hippocampal formation. To examine a more general long-term memory enhancing effect of nuclear ERK potentiation, we next injected RB5 directly into the dorsal hippocampus of rats before contextual fear conditioning, a learning task in which allows to dissociate between short-term and long-term memory ^24^. A single bilateral infusion of RB5 20 min into the dorsal hippocampus prior to conditioning markedly enhanced fear memory acquisition (post US), short-term memory (3h) and long-term memory (48h) (Fig. 4d). These data confirm that a localized infusion of a selective nuclear ERK enhancing drug improves hippocampal-dependent learning and memory.

Mild cognitive impairment (MCI) is a prodromal stage observed across most neurodegenerative disorders including AD and HD. The ability of RB5 to significantly enhance memory formation and consolidation could be exploited to prevent MCI in models of such brain disorders. Thus, we first investigated whether a single systemic RB5 injection could ameliorate the cognitive impairment observed in the Tg2576 mouse, which has previously been validated as a MCI model of AD ^25^. As shown in Fig. 4e, a single injection of RB5 fully prevented the memory impairment observed in contextual fear conditioning in the Tg2576 mouse. This confirms the potential therapeutic validity of a selective enhancement of nuclear ERK using this early model of AD.

To test the possibility that RB5 could ameliorate procedural learning deficits in a genetic model of HD, we used the relatively early onset zQ175 mouse model, containing a CAG expansion in Htt of approximately 180 repeats ^26,27^. Animals were trained in a procedural task (‘nose poking’) using 9-hole operant boxes, as previously described for the Hdh^Q111/+^ knock-in mouse ^28^. Mice were trained to nose poke on a simple fixed ratio (FR1) schedule of reinforcement. Once trained, male and female animals were subdivided and injected with either RB5 (10mg/kg, i.p.) or scramble peptide 1h before being tested daily on FR1 schedule (Fig. 4f). After 10 days, both male and female zQ175 showed clear deficits in nose poke responses as expected. However, upon RB5 administration a clear sex difference was observed. Both wild type (WT) and zQ175 female mice showed a significant enhancement of the responses over 9 days of RB5 treatment (from day 11 to day 19). Remarkably, the RB5 treated zQ175 female mice reached the same level of performance as the scramble treated WT mice, confirming that RB5 is effective in restoring the cognitive impairments in zQ175 model of HD. In marked contrast, RB5 treatment did not alter the responses in males of zQ175 strain, independently from genotype (data not shown).

In sum, this experiment indicates that RB5 acts as cognitive enhancer in females trained in a striatal-dependent procedural learning task and rescued procedural learning deficits in zQ175 HD mice. These data are in accordance with previous evidence showing that early experiences of environmental enrichment with cognitive testing not only ameliorated the severity of cognitive deficits in zQ175 mice but this beneficial effect was mostly evident in female mice ^29^. At present, the sex difference is not fully understood and will require further investigation.

Finally, we explored the possibility of a similar cognitive enhancing effect also in another genetic HD mouse model. Hdh^Q111/+^ is a late onset disease model, manifesting cognitive decline after 14-18 months of age and a moderate neurodegeneration ^28,30^. Since we wanted to maximize the therapeutic effect by treating animals as early as possible, we injected a LV expressing ERK2 bilaterally into the dorsal striatum at 2 months of age to sustain ERK signalling throughout this longitudinal experiment. Mice were then behaviourally tested at 18 months of age. We trained all four groups in an operant FR1 schedule of reinforcement and data indicated that three groups (WT-GFP, WT-ERK and Hdh^Q111/^+ -ERK2) learned the task while the Hdh^Q111/^+ -GFP group did not (Fig. 4g).

Altogether, these data indicate that a pharmacogenetic potentiation of ERK signalling in the brain can prevent MCI and the cognitive decline measured in distinct mouse models of neurodegenerative disorders.

## Discussion

In patients affected by AD, PD or HD the molecular processes underlying disease may take decades to manifest in the earliest behavioural symptoms and occur in the absence of major anatomical alterations and neuronal cell loss. It is not disputed that early intervention is the key for successful treatments and the overarching goal is to prevent synaptic failure associated with cognitive deterioration. This is particularly evident for dementia research where the outcome of recent trials aiming at preventing αβ amyloid accumulation in AD that have not been matched by a corresponding clinically relevant attenuation of the cognitive symptoms ^31^. Potential interventions during the phase when prodromal cognitive symptoms appear, such as mild cognitive impairment (MCI) has not yet been fully exploited ^32^.

Here we have provided initial evidence for a strategy targeting a common mechanism implicated both in the control of cell survival and in cognitive enhancement early in the development of neurodegenerative disorders such as HD and AD. Since the discovery of its role in learning and memory ^33,34^, the Ras-ERK pathway has been subjected to intense scrutiny as a therapeutic target for both psychiatric and neurological disorders, largely through the use of MEK inhibitors and in a few cases, through proof-of-principle genetic manipulations in the mouse ^3^. Both approaches were associated with significant limitations and in some cases generated conflicting results. On one hand, the use of MEK inhibitors can only demonstrate a permissive for the pathway role (“is ERK signalling necessary?”) but precludes any investigation of the potential instructive role (“is ERK signalling sufficient?”). On the other hand, genetic models selectively alter the expression of a key component of the pathway either early in development (global gene manipulation) or for a prolonged period of time (weeks to months, in case of tissue/cell specific gene targeting) whose effects maybe confounded by associated cellular adaptations and compensatory mechanisms ^35,36^.

The same limitations also apply to the dissection of the other important role of ERK signalling relevant to therapeutic intervention, i.e. the control of neuronal cell survival and cell death. In the last decade, contrary to reports indicating that the ERK cascade is essentially a pro-survival pathway, circumstantial evidence, mainly in cultured cells, indicate that this signalling pathway may also play a pro-apoptotic (or pro-cell death in general) role ^37,38^.

In order to circumvent these limitations, we used two complementary approaches in the adult, selective ERK1 or ERK2 gene inactivation and a novel nuclear ERK enhancing cell penetrating peptide. Both approaches allow manipulations of the system that were precluded in the previous studies. First of all, by applying selective ERK1 and ERK2 gene knockdowns using viral vectors we achieved a rapid removal of the two proteins independently from the other in the brain *in vivo*, without the associated developmental effects. Secondly, the cell penetrating peptide allowed us to determine for the first time whether a selective nuclear ERK enhancement could impact *in vivo* on cell survival, synaptic plasticity and memory.

The present research originates by the observation that ERK1 KO mice showed a potentiation of ERK2-mediated signalling which resulted in enhanced neuronal plasticity and memory ^4,5^. We first confirmed and extended our original findings that a bidirectional modulation of the ERK2/ERK1 ratio could impact on global ERK signalling and alter cell survival in normal mice and in models of neurodegeneration. Most importantly, a selective activation of nuclear ERK signalling not only enhanced memory in wild type animals (both rats and mice), but also rescued MCI in an AD mouse model and ameliorated cognitive deterioration in two distinct genetic models of HD. We believe that the effect on memory is a primary phenomenon, occurring independently from the attenuation of neuronal cell loss seen in models of AD, HD and PD. However, the observation that a sustained ERK2 overexpression in the striatum prevents memory decline and also significantly attenuates caspase 3 staining strongly suggest that ERK signalling potentiation has a dual beneficial long-term effect, at least in HD models. This is consistent with a well-established role for ERK downstream to BDNF-TrkB receptor activation in the striatum ^37,39^. BDNF (or therapeutics modulating its action) administration to HD patients has not yet been successful ^40^. In principle an approach using RB5 seems feasible, but will require further investigation. The observation that nuclear ERK modulation ameliorates cognitive deficits in HD models as well as in WT animals suggests an interesting hypothesis. It has been previously shown in a number of HD models, including HdhQ111 and zQ175, that behavioural training or environmental enrichment (EE) can lead to a significant delay in cognitive decline ^29,41–43^. Interestingly, these effects in HD mouse models have been shown to be gender biased, with females better responding to the treatment, and were paralleled by the alleviation of depression-like behaviour normally more pronounced in females than in males ^44^. While the gender effect observed across HD mouse models has not yet been fully confirmed in HD patients, the “therapeutic” effect associated with behavioural training/EE is consistent with a prominent role of epigenetic mechanisms in overcoming the deleterious expression of mutant huntingtin. The ability of our peptide to mimic the same epigenetic mechanisms provides an unprecedented opportunity for developing novel pharmacological approaches for Huntington’s disease, as well as for the cognitive enhancement of the healthy population.

Much less is known of the role of brain ERK signalling in AD. Early reports indicate that ERK signalling components, including ERK1/2 and MEK1/2, are upregulated in post mortem AD and other dementias brains ^45–47^. These findings have led to the speculation that ERK signalling may contribute to the pathogenic process in a complex mechanism by which αβ oligomers can activate ERK, leading to tau phosphorylation and further αβ amyloid accumulation ^48–50^. However, recent evidence indicates that ERK does not phosphorylate tau in vivo ^51^. Moreover, intrahippocampal injection of αβ oligomers rapidly downregulates ERK phosphorylation, thus correlating with observed memory impairments ^52^. Although early evidence suggested that ERK2 activity in the Tg2576 mouse model of AD was only attenuated at 20 months of age ^53^, here we showed a significantly diminished ERK phosphorylation at 7 months of age that could be rescued by the ERK enhancing treatment. Our interpretation of these data is that ERK signalling is likely to response to αβ amyloid toxicity as a compensatory mechanism, rather than being part of the pathogenic process.

Altogether, we have unravelled a novel mechanism that may provide both neuroprotection and cognitive enhancement across neurodegenerative diseases. In addition, we have also demonstrated that a selective potentiation of nuclear ERK signalling in the brain might lead to a significant cognitive improvement in normal animals, with a potentially interesting perspective for delaying mental deterioration in the general ageing population.

## Materials and Methods

### Animals

All experiments were conducted according to the European Community guide-lines (Directive 2010/63/EU) and previously approved by the Institutional Animal Care and Use Committee (IACUC) protocols (# 477) of the IRCCS-San Raffaele Scientific Institute and UK Home Office project license to RB (PPL 30/3036) at Cardiff University.

CD1 male mice (Charles River), 2 months old were used for immunohistochemical analysis (time course, dose-response, mass spectrometry) and Novel Object Recognition test. C57BL/6 male mice (Charles River) 2 months old were used for LV injections, ex vivo analysis, IHC and IF. C57BL/6 male mice of 3 months were used for MPTP intoxication. ERK1 KO male and littermate controls, generated as previously described ^4^ were used for ex vivo analysis. Heterozygous Hdh^Q111/^+ knock-in mice (Jax1, Bar Harbour, Maine, U.S.A.) were bred in-house on a C57BL/6J background. Heterozygous zQ175 knock-in mice carrying approximately 180 CAG repeats (CHDI-81003003, Psychogenics, Inc. Tarrytown, NY) of 5-7 months and Tg2576 transgenic male mice (C57BL6 and SLJ mix background, Taconic Biosciences) between 5 and 7 months were used for IHC, IF and CFC test. Lister Hooded male rats (280–350gr) were purchased from Harlan Laboratories and used for CFC.

### Lentiviral vectors production

Generation and production of Lentiviral (LV) constructs for RNAi of ERK1 and ERK2 (both in shRNA and mir RNA configuration) as well as LV overexpressing ERK1, ERK2 and their swapped counterparts have been previously described ^8,54^. Briefly, third generation non-replicative LVs were modifications of the originally described backbone ^55^. LVs have been prepared in HEK 293T cells with a final titre > 10^9^ T.U. (transforming units). Cells and mice have been transduced at multiplicity of infection (moi) >5.

### Mouse embryonic fibroblasts (MEF) preparation and LV infection

MEF cultures were prepared from wild type E13.5 embryos obtained from C57Bl/6 mice as previously described ^54^. Then MEF were fixed 3 or 5 days after LV infection in PFA 4% for 25 minutes at RT, rinsed in PBS 1X and then stored in 70% ethanol at −20°C.

### Neuronal cultures and LV infection

Neuronal cultures were prepared as previously described ^4^. Briefly, embryonic cultures (E17) were prepared from cortices of WT mice, plated onto poly-L-lysine coated glass and kept for 10 days in culture medium. On day 3 cells were infected with LVs of interest (moi=5) and 3, 5 or 7 day later were fixed in PFA 4% for 25 minutes at RT, rinsed in PBS 1X and stored in 70% ethanol at −20°C.

### LVs in vivo injections

C5BL/6 adult male mice were deeply anaesthetized (i.p. 8% Ketamine (Ketavet 100, Intervet) and 4% Xylazine (Rompun 20mg/ml, Bayer) and secured on a stereotaxic frame (Kopf). Two bilateral LV injections (2ul each) were performed in the dorsal striatum at the following coordinates: site 1: AP +1; L −2.1; DV −2.6; site 2: AP +0.3; L −2.3; DV −2.4. LV of interest was injected either in right or left hemisphere according to a randomised design as control LVs. Two weeks post-injections, mice were anaesthetized and perfused with ice-cold buffered 4% PFA. Brains were extracted, post-fixed overnight and transferred to 30% buffered sucrose for 24 hours. Coronal sections were cut to a 35 μm thickness on a freezing microtome and stored in a cryoprotective solution at −20°C until processing for IHC or IF.

2 months old Hdh^Q111/+^ knock-in mice were deeply anaesthetized with Isoflurane (Piramal Critical Care) and secured on a stereotaxic frame (Stoelting). Two bilateral LV injections (1ul each) were performed in the dorsal striatum at the following coordinates: site 1: AP+1.2,L-1.4, DV-3, site 2: AP +0.3, L −2.3, DV −2.4, 16 months post-injections, mice were subjected to behavioural tests.

### TUNEL Assay

Detection of apoptosis on cultured cells or tissue sections was carried out using the DeadEnd™ Colorimetric TUNEL System kit (Promega). Briefly, slides were immersed in PBS for 5 minutes RT and then incubated for 25 minutes in 20μg/ml Proteinase K solution. Then, the slides were rinsed in PBS 1X, re-fixed in PFA 4% for 5 minutes and rinsed twice with PBS 1X. 100μl of equilibration buffer was added to each tissue section for 5-10 minutes followed by 1h of incubation at 37°C in a humidified chamber with 100μl of rTdT reaction mix. After that, slides were immersed in 2X SSC solution in a Coplin jar for 15 minutes and washed twice in PBS 1X for 5 minutes. The endogenous peroxidase activity was blocked with 0.3% hydrogen peroxide for 5 minutes. The slides were rinsed twice in PBS 1X for 5 minutes and incubated for 30 minutes with Streptavidin HRP solution and finally DAB substrate was added to react for 10 min. The slices were developed using DAB, and washed several times using deionized water. Pictures from all experimental groups were acquired through a light microscope.

### Imaging of dendritic spines

Mice injected with LVs were let expressing the transgene for four weeks. Perfused brains stored in PFA 4% were cut at the vibratome at 250-30 μm thickness. Slices were mounted on slides. The measure of the dendritic spine density was carried out using a two-photon microscope (Ultima IV, Prairie Technology) equipped with a 10W laser (Chamaleon Ultra 2, Coherent) tuned at 890 nm that delivered about 30mW at the sample. Imaging was restricted to well resolved dendrites avoiding fields that were too cluttered. Images were acquired with a water immersion lens (Olympus, 60, numerical aperture: 0.95) at a resolution of 512×512 pixels with zoom 4 leading to a field of 50.7×50.7 mm and a nominal linear resolution of about 0. 1 mm per pixel. Selected dendrites were acquired by collecting a Z-stack with a vertical resolution of 0.75 um. Images were analyzed with ImageJ.

### Peptide synthesis

A peptide for inhibiting ERK1 (*Mapk3*) protein kinase signalling was designed around the N-terminal domain of ERK1 and hereafter named RB5. RB5 peptide and its ineffective Scramble version contain a TAT sequence able to translocate the plasma membrane.

RB5 (GRKKRRQRRRP<u>PQ</u>GGGGGEP<u>RR</u>TEGVGPGVPG<u>E</u>V<u>E</u>MVKGQPFD<u>V</u>) and SCRAMBLE (GRKKRRQRRRPPRVGPGVPEGVGVAVFGVKEPGQTGDVGPVGE) peptides were custom synthesised by GENECUST EUROPE (Luxembourg).

For all *in vitro* and *in vivo* experiments batches of 200 mg, highly purified by high-performance liquid chromatography (HPLC) (≥ 95 %) were used. In order to enhance stability, RB5 was synthesised containing six D-aminoacid isomers in substitution of the native L-aminoacids (underlined including the C-terminal aminoacid) and the acetylated N-Terminal aminoacid. RB5 and Scramble peptides were dissolved in PBS1X and injected at the appropriate concentrations.

### Liquid chromatography and tandem mass spectrometry (HPLC-MS/MS)

Bioavailability of RB5 in the mouse brain was determined at different time intervals (1, 3, 6,12 h) after a single i.p. adminstration of 20 mg/kg. RB5 was then quantified in mouse brain using HPLC-MS/MS. Brain samples were homogenized with 1:4 w/v of 50% acetonitrile, 5 % TFA in water with an homogenizer ultra-turrax and then centrifuged at 13000 rpm for 10 min at 4°C. The supernatant was collected and centrifuged at 13000 rpm for 2 minutes at 4°C. The supernatant was then collected in ice, extracted using Sep-Pak cartridges C18 lyophilised and kept at 4°C before HPLC-MS/MS analysis. Immediately before the analysis samples were suspended in 100 μL of 0.1% HCOOH in water/8% acetonitrile in auto-sampler vials.

HPLC-MS/MS analysis was performed using a system consisting of an Agilent 1200 series HPLC system coupled with an Agilent 6410 Triple Quadruple mass spectrometer. Mass Hunter Workstation v. B.01.03 software was used for data collection and processing (Agilent Technologies, Santa Clara, California, US). The quantification of mouse brain levels of RB5 and scramble peptides was carried out using an internal standard curve with peptide concentrations ranging from 0.1 to 4 ng per μl. RB5 and scramble peptides and the internal standard were separated at room temperature by injecting 10 μL of extracted sample onto a Jupiter C4 300 A analytical column, 2×150mm, 5μm particle size (Phenomenex, CA). Gradient elution was used for chromatographic separation, using 0.1% formic acid in water as solvent A, and acetonitrile as solvent B at a flow rate of 200 μl/min. The elution started with 92% of eluent A and 8% of eluent B maintained for 1 min, followed by a 4 min linear gradient to 75% of eluent B, a 1 min linear gradient to 99% of eluent B, a 2 min isocratic elution and a 0. 5 min linear gradient to 8% of eluent B, which was maintained for 9.5 min to equilibrate the column. The samples were maintained at 4 °C in the autosampler.

Peptides were then detected on an Agilent 6410 QQQ mass spectrometer using the following parameters: positive ion mode, 5 kV capillary voltage, cone voltage 500 V, gas flow rate 8 L/min at 350 °C, nebulizer gas pressure 40 PSI at 350 °C, well time 75 msec and Q1 and Q3 set to unit resolution.

### Ex-vivo system in acute brain slices

Adult C57Bl/6 mice were decapitated after cervical dislocation and brain slices were freshly prepared as previously described in ^8,56^. Briefly, the brains were rapidly removed from the skull, submerged in ice-cold sucrose-based dissecting solution and sliced into 200 *μ*m-thick slices using a vibratome. Striatal slices were subsequently transferred into the brain slice chamber and allowed to recover for 1h at 32°C, with a constant perfusion of carboxygenated artificial cerebrospinal fluid in the presence of RB5 peptide (0.5 – 100μM for IC50 determination). RB5 or scramble inactive peptide (50μM) were used in combination with 100μM glutamate to stimulate brain slices into the chamber for 10 min. After a rapid fixation in 4% PFA, slices were rinsed and cryoprotected overnight at 4°C in sucrose solution 30%. On the following day, slices were cut into 18 *μ*m-thick slices using a cryostat (Leica CM1850), mounted onto SuperFrost Plus slides (Thermo Scientific) and stored at −20ºC until processing for immunohistochemistry or immunofluorescence.

### In vivo administration of drugs for histochemical analysis

3-NP (3-Nitropropionic acid) (N5636, Sigma) was dissolved in distilled water to a concentration of 50 mg/ml (pH 7.4) and passed through a 0.2-μm filter and kept at −80°C until use. The 3-NP solution was administered i.p. once daily at a dose of 50 mg/kg either for 21 consecutive days in the LV injected mice or for 7 days in combination with RB5.

1-methyl-4-phenyl-1,2,3,6-tetrahydropyridine hydrochloride (MPTP-HCl, Sigma, Italy) was dissolved in saline and administered i.p. once daily at the dose of 20 mg/kg for 4 days.

Tg2576 mice were administered either with RB5 or scramble (20mg/kg i.p.) for 7 days for IHC analysis.

### Immunohistochemistry

Coronal sections from the SNc (40-μm thick) of MPTP-treated mice were cut on a vibratome. Immunohistochemistry was performed as described in ^56^. Briefly, 1h after blocking in 5% normal goat serum and 0.1% Triton X-100 solution, free-floating sections were incubated overnight at 4°C with one of the following primary antibodies: anti-phospho-p44/42 MAP kinase (Thr202/Tyr204) (1:200, Cell Signalling Technology #9101), anti-phospho–Elk-1 (Ser383) (1:100, Cell Signalling #9181), anti-phospho-Kv4.2 (Thr607)-R (1:100, Santa Cruz sc22254-R), anti-phospho-MSK-1 (1:100 Cell Signalling Technology #9595), anti-phospho-MEK1/2 (Ser218/222) (1:100, sc-7995 Santa Cruz Biotechnology), c-Fos (1:100, sc-52 Santa Cruz Biotechnology) and tyrosine Hydroxylase (monoclonal 1:1000, Sigma Aldrich, Italy) overnight at 4°C Sections were then incubated with biotinylated goat anti-rabbit IgG (1:200, Vector Laboratories) for 2h. the classic avidin–peroxidase complex (ABC, Vector, UK) protocol was applied, using 3,3’-diaminobenzidine (DAB, Sigma) as a chromogen.

### Immunofluorescence

Immunofluorescence was performed as described in ^23,56^. Briefly, 1h after blocking in 5% normal goat serum, slices were incubated overnight at 4°C with the following primary antibodies: anti-phospho (Ser10)-acetylated (Lys14) histone H3 (1:1000, Millipore), anti-phospho-S6 ribosomal protein (Ser235/236) (1:200, Cell Signalling Technology) anti-phospho-S6 ribosomal protein (Ser240/244) (1:200, Cell Signalling) or Cleaved Caspase 3 (1:200, Cell Signalling Technology) and NeuN (1:1000, Millipore). Corresponding secondary antibodies (Alexa Fluor 546 conjugated anti-mouse (1:200, Thermo Fisher Scientific), Alexa Fluor 488 conjugated anti-rabbit (1:500, Thermo Fisher Scientific) and Pacific Blue goat anti-mouse (1:200, Thermo Fisher Scientific) were incubated at room temperature for 1h.

Single and double-labelled images (1024 × 1024 *μ*m) were obtained at 40X magnification from striatum using a laser scanning confocal microscopy (Leica SP2) equipped with the corresponding lasers and appropriate filters sets to avoid the cross-talk between the fluorochromes.

### IHC and IF Image quantification

For *ex-vivo* immunohistochemistry experiments, images were acquired from the dorsal striatum using a bright-field microscope (Macro/Micro Imaging System, Leica) under a 40X magnification. Neuronal quantification was performed with ImageJ software by counting the number of phospho-ERK positive cells in 2 sections for each sample in 3 fields per section. The number of positive cells per mm^2^ in RB5-treated slices was normalized over the scramble controls. The EC50 for pERK was calculated using GraphPad Prism software.

For *in vivo* immunohistochemistry experiments, images were acquired from the dorsal striatum using a Leica DM IRB microscope under a 20X objective. The number of p-ERK, p-MSK-1, p-ELK, p-MEK-1, p-VG K+, and c-Fos positive cells was counted in 2-3 consecutive sections per mouse, bilaterally and averaged across the sections using the ImageJ software by a blind investigator to condition. Estimation of nigral cell loss was assessed by quantifying the number of TH-immunopositive neurons of the SNc in 360 μm-spaced sections for each animal.

In *ex-vivo* and *in-vivo* immunofluorescence experiments, the number of p-S6 or p-AcH3 immunoreactive neurons among NeuN positive neurons was counted in 2-3 consecutive striatal sections per mouse in 4 fields per section. Caspase 3 positive cells were quantified among NeuN positive neurons in each slide and expressed as percentage (ratio between Caspase3 positive cells and total NeuN positive neurons) for each field acquired through the slide.

### Spontaneous rotations

Mice were unilaterally injected in the striatum with LVs of interest (2 sites, 2ul each) as above described. The contralateral sides were injected with control LV to minimise the effect of surgery. 2 weeks later, spontaneous rotational behaviour was observed in a square arena and the number of 180^0^ rotations either side were measured. Positive values represent net ipsilateral rotations while negative values represent net contralateral rotations (the number of 180^0^ rotations towards LV injected side - number of 180^0^ rotations towards control side).

### Novel Object Recognition Test (NOR)

NOR was performed as previously described ^22,23,56^. Briefly, the test was performed in an open square box placed in a quiet room with dim light. The objects used were: parallelepipeds in metal and glass vials filled with water. The protocol required three days and it was performed as follows:

Day 1: mice were individually placed in the empty arena for 5 min in order to familiarize with it and to measure their anxiety (thigmotaxis trial). The percentage of thigmotaxis is calculated as the time spent in the peripheral zone out of the total time spent in the arena (300 sec.). Animals showing a thigmotaxis > 90% are discarded from the sample because considered biased for anxiety. Day 2: mice were placed into the arena for 10 minutes, where they were allowed to explore two identical objects (training trial). The amount of time spent by mice exploring each object was scored. Day 3: one of the two identical objects (parallelepiped) was changed with a new object (glass vial) and the mice were allowed to explore them for 10 minutes (test trial). Times of exploration for the familiar object and for the novel object were recorded. The discrimination index (DI), defined as the difference between the exploration time for the novel object and the one for the familiar object, divided by total exploration time, was calculated. The sessions were recorded with the video tracking software SMART (Panlab, Barcelona, Spain).

RB5 or Scramble peptides (20mg/kg, i.p.) were administered 1h before the training session on day 2 and then long term memory was tested in independent groups at different time points: 24, 48, 72, 120 and 168 hours post injection.

### Contextual Fear Conditioning Test (CFC)

Lister Hooded male rats (280–350gr) were housed in pairs, in holding rooms maintained at 21°C on a reversed-light cycle (12 h light/dark; lights on at 10:00 P.M.). All experiments were conducted in the dark period of the rats. RB5 and Scramble peptides were infused into dorsal hippocampus.

#### Surgery and Microinfusions into the Dorsal Hippocampus

Steel double guide cannulae aimed at the dorsal hippocampus (AP −3.50, relative to bregma) were surgical implanted under anaesthesia at least one week prior to behavioural training and microinfusions. Bilateral infusions with either 2 mg/ml Scramble or RB5, 20 min prior to conditioning (pH 7.0, 1.0 ml/side, rate = 0.5 ml/min) via the chronically indwelling cannula were carried out in awake rats using a syringe pump, connected to injectors (28 gauge, projecting 1 mm beyond the guide cannulae) by polyethylene tubing.

#### CFC rat protocol

Conditioning was performed in one of two distinct contexts. These contexts were designed to differ in a number of distinctive characteristics including size, spatial location, odour and lighting. During the 3 min conditioning training trial, rats received a single scramble footshock (0.5 mA for 2 s) 2 min after being placed into one of the conditioning contexts (CtxA). All rats were returned to the home cages after conditioning. Retrieval tests 3h (post-retrieval short-term memory, STM), or 2 days (long-term memory, LTM) after recall again consisted of exposing the rat to the conditioning context for 2 min not delivering a foot shock.

#### CFC mice protocol

7 months old Tg2576 mice and WT mice were treated with either Scramble or RB5 (20 mg/kg, i.p.) 1h before CFC. Mice were individually placed in the conditioning chamber for 120 s of free exploration followed by five foot shocks (0.7 mA, 2-s duration, separated by 60-s intervals) delivered through the grid floor. Context Fear Memory was assessed 24 h later by returning mice for 5 min to the conditioning chamber and not delivering a foot shock.

For all protocols, freezing behavior served as a measure of conditioned fear to the context during the conditioning and retrieval tests.

### 9-hole operant boxes test

Operant testing was conducted in 16 9-hole operant boxes as previously described ^28^. Each operant box contains a horizontal array of nine holes with infrared beams localised to the front of each hole to detect nose pokes. A peristaltic pump delivers liquid reinforcement (strawberry milk) into a magazine at the front of the box.

A week before starting the training, mice undergo water restriction for 18 hours/day and are kept under this regimen throughout the experimental procedures. Mice are taught to nose poke on a simple fixed ratio (FR1) schedule of reinforcement: to obtain reward, mice are required to respond to a stimulus light in the central hole via a single nose poke. zQ175 mice were trained daily on this program during 20 minutes sessions for 10 days (training phase). Once trained, animals were subdivided in 4 groups and injected with RB5 or scramble (10mg/kg, i.p.) peptides 1h before being tested on FR1 schedule for other 9 days.

### Electrophysiology (Hippocampus)

#### Slice preparations

Acute transverse hippocampal slices (400 μm thick) were prepared from adult male Sprague-Dawley rats (p60-p90) after a lethal dose of isoflurane inhalation in accordance with the Home Office guidelines and a directed by the Home Office Licensing Team at Cardiff University, as previously described ^57^. The hippocampi were dissected in ice-cold slicing solution containing (in mM) 110 choline chloride, 25 glucose, 25 NaHCO_3_, 2.5 KCl, 1.25 NaH_2_PO_4_, 0.5 CaCl_2_, 7 MgCl_2_, 11.6 Na ascorbate, and 3.1 Na pyruvate, then mounted on agar and cut using a Microm HM 650 V vibratome (Thermo Scientific). Slices were incubated in artificial cerebrospinal fluid (aCSF) containing (in mM) 119 NaCl, 10 glucose, 26.1 NaHCO_3_, 2.5 KCl, 1, NaH_2_PO_4_, 2.5 CaCl_2_, and 1.3 MgCl_2_ at 36 °C for 30 min, the stored at room temperature until use. All solutions were equilibrated with 95% CO_2_ and 5% O_2_ and had osmolarity of 300-310 mOsm. Slices were then cut between CA3 and CA1 before being transferred to the recording chamber.

#### Patch-clamp recordings

Whole-cell patch clamp recordings were made form CA1 pyramidal neurons visualized under differential interference contrast (Olympus BX51 WI) in a submerged recording chamber perfused with aCSF (~ 2 ml/min) at 35 °C supplemented with 50 μM picrotoxin to block GABA_A_ receptors. Patch electrodes (3-5 MΩ) were pulled from borosilicate filamented glass capillaries on a P-1000 Flaming-Brown micropipette puller (Sutter Instruments) and filled with intracellular solution (in mM): 117 KMeSO 3, 8 NaCl, 1 MgCl 2, 10 HEPES, 0.2 EGTA, 4 MgATP and 0.3 Na2GTP, pH 7.2, 280 mOsm. For experiments using RB peptides, the intracellular solution was supplemented freshly with either RB5 or control peptide (Scr) at a final concentration of 50 μM and the pH was readjusted with 1N KOH. Cells were voltage-clamped at −70 mV. Recordings were made with a Multiclamp 700B amplifier (Molecular Probes) analog filtered at 10 kHz and digitized at 50 kHz using a Axon Digidata 1550 acquisition system and Clampex software. Recordings were initiated immediately after establishing whole-cell configuration. Synaptic responses were evoked by delivering 0.1-1 ms square pulses at 0.1 Hz through a bipolar tungsten electrode inserted in *stratum radiatum* (Digitimer Constant Current Stimulus Isolation unit). Consecutive EPSCs were averaged offline every minute and the amplitude was normalized to the average of first minute of recording. Series resistance (Rs) was monitored throughout and cells with Rs > 30 MΩ or with over 20% change in Rs were discarded.

### Electrophysiology (Striatum)

#### Slice preparation

Brain slices containing both the striatum and the cortex were prepared as described ^20^. Mice were anesthetized by isofluorane and decapitated, and their brains were rapidly transferred to ice-cold dissecting aCSF containing (in mM): 110 Choline-Cl, 2.5 KCl, 1.25 NaH_2_PO_4_, 7 MgCl_2_6H_2_O, 0.5 CaCl_2_, 25 NaHCO_3_, 25 D-glucose, 11.6 ascorbic acid, saturated with 95% O_2_ and 5% CO_2_. Horizontal corticostriatal slices (270μm thick; patch-clamp recordings) were cut in the dissecting aCSF using a Vibrotome 1000S slicer (Leica, Italy), then transferred to normal aCSF containing (in mM): 115 NaCl, 3.5 KCl, 1.2 NaH_2_PO_4_, 1.3 MgCl_2_6H_2_O, 2 CaCl_2_, 25 NaHCO_3_, and 25 D-glucose and aerated with 95% O_2_ and 5% CO_2_. Following 20 minutes of incubation at 32°C, slices were kept at RT. During experiments, slices were continuously perfused with aCFS at a rate of 2 ml/min at 28°C.

#### Identification of dSPNs and iSPNs

To identify SPNs of the direct (dSPN) and indirect (iSPN) pathways, neurons were filled with Neurobiotin (0.5 mg/ml, DBA Italia, Segrate, Italy) during recordings and subsequently processed for immunostaining of the A2A receptor (marker of iSPNs) and substance P (marker of dSPNs) ^20^. After recording, slices were fixed with 4% PFA in 0.1 M PB (pH 7.4) overnight at 4°C and then incubated in primary antibodies. Rabbit polyclonal antibody to A2A (1:250, Enzo Life Sciences, Farmingdale, New York) and rat monoclonal antibody to substance P (1:200, Millipore, Billerica, Massachusetts) were diluted in 0.1 M PB containing 0.3% Triton X-100. Sections were subsequently incubated with Alexa 647- or Alexa 488-conjugated secondary antibodies (1:200) and Alexa 568-conjugated streptavidin (1:1000) (Invitrogen, Carlsbad, California), mounted on glass slides, and coverslipped. Images were acquired with an inverted Leica TCS SP5 AOBS TANDEM confocal microscope.

#### Patch-clamp recordings

Whole-cell recordings were made under direct IR-DIC (infrared-differential interference contrast) visualization of neurons in the dorsolateral striatum. Current clamp experiments were performed by using borosilicate patch pipettes (4-6 MΩ) filled with a potassium-methylsulfate-based internal solution containing (in mM): 135 KMeSO_4_, 10 KCl, 10 HEPES, 1 MgCl_2_, 2 Na_2_-ATP, 0.4 Na_3_-GTP (pH 7.2-7.3, 280-290 mOsm/kg), completed, at the beginning of each experimental day, with RB5 peptide or its scramble-peptide control which were diluted at the final concentration of 50μM. Since, each peptide was dissolved in acetic acid to have 2.1mM stock, the pH of the potassium-methylsulfate-based internal solution was adjusted with NaOH every time after adding peptides.

SPNs were clamped at a holding membrane potential of −80mV. Excitatory postsynaptic potentials (EPSPs) were evoked in the presence of the GABA_A_ receptor antagonist gabazine (10 μM) by cortical stimulation from the somatosensory cortex layer 5 by using a concentric bipolar electrode (80μsc-200μsc, 0.9mA-1.6mA, CBAPB75, FHC, Bowdoin, ME) connected to a constant-current isolation unit (Digitimer LTD, Model DS3) and acquired every 10 seconds. During HFS-plasticity induction (4 × 1 s long 100-Hz trains, repeated every 10 s), the postsynaptic cell was depolarized from −80 to −50 mV. Signals were sampled at 20kHz filtered to 10kHz. During the experiments, only cells with a stable resting membrane potential ≤ −78 mV were included in the analysis. Series resistance (range 15-25 MΩ) was monitored at regular intervals throughout the recording and presented minimal variations (≤20%) in the analysed cells. Data are reported without corrections for liquid junction potentials. Data were acquired using a Multiclamp 700B amplifier controlled by pClamp 10 software (Molecular Device), with a Digidata 1322 (Molecular Device).

#### Data Analysis

The occurrence and magnitude of synaptic plasticity was evaluated by comparing the normalized EPSP amplitudes from the last 5 min of baseline recordings with the corresponding values at 20–30 min after HFS (HFS-LTD). LTD plots were generated by averaging the peak amplitude of individual EPSPs in 1-min bins. Data were analysed by One-way repeated-measures ANOVA (RM1W) for comparisons within a group. Post hoc analyses (Tukey, multiple comparison tests) were only performed for ANOVAs that yielded significant main effects. Two groups were tested for statistical significance using the two-population t-test and Mann-Whitney U-nonparametric test (GraphPad Prism 6 software).

## Acknowledgments

This work was funded by the Italian Ministry of Health, the Compagnia di San Paolo, the Welsh Government (Life Science Bridging Fund) (to RB), the Medical Research Council (to SF), the Waterloo Foundation “Changing Minds” Programme and the Wellcome Strategic Award “DEFINE” (to JH), the Fondazione Telethon (grant no. GGP15098), Albero di Greta ONLUS, International Foundation for CDKL5 Research, Lou Lou Foundation and Fondazione CRT (to MG).

